# Synchronization of iPS-derived cardiomyocytes to visitor heartbeat in an interactive museum exhibit

**DOI:** 10.1101/2021.04.19.440451

**Authors:** Juan A. Perez-Bermejo, Samuel J. Reisman, Joyce Ma, Chris Cerrito, Bruce R. Conklin, Kristina Yu

## Abstract

Science museums play an important role in science education, both engaging the public with science concepts and building support for scientific research. Designing museum exhibits to meet increasing public interests in the life sciences is particularly important, yet remains challenging. In this report we describe *Give Heart Cells a Beat*, a permanent interactive science exhibit that allows museum visitors to synchronize the beating of live stem cell-derived cardiomyocytes to their own heart rate in real-time. Evaluation with museum visitors reveals that the exhibit engaged the public with the specimen and prompted curiosity in heart biology and, to a lesser degree, stem cells and electrophysiology. *Give Heart Cells a Beat* is the product of a close collaboration between a museum and a research laboratory, and, to our knowledge, the first example of the use of live human heart cells in an interactive exhibit. We hope this exhibit will serve as an example for the implementation of stem cell technology in the field of informal science education and encourage others to pursue close working relationships between academia and public science venues such as museums.

## Introduction

Important cell biology advances, such as the development of stem cell technologies, have led to breakthroughs in basic and translational biology^1^ and are increasingly relevant to public life^2–5^. This scientific and societal prominence has created an opportunity for scientists and educators to bring these technologies to the public in an engaging and relatable way, and to use them to create new types of educational experiences. Museums and science centers are key components of the science education landscape^6–8^, and provide rich opportunities for reaching a wide and diverse audience^6^. In addition, there is a growing appreciation for interactive museum exhibits^6,9–11^ and the use of real living samples, rather than recordings or simulations^11^, to promote visitors’ interest, engagement and understanding of the content. However, the development of interactive biology exhibits featuring live human cells has been significantly limited by cost and availability of reagents, capabilities needed for museums to maintain and display cell lines, and lack of mechanisms that let visitors interact with microscopic specimens. As a result, human cell biology at science museums has traditionally been limited to non-interactive models, fixed samples, or simulations. Academic research laboratories have been called on to expand their involvement in science education^12^ and represent a promising partner in overcoming these challenges^13^.

Here we report our development and assessment of *Give Heart Cells a Beat* (*GHCB*), an exhibit that allows visitors to interact with living human induced pluripotent stem (iPS) cell-derived cardiomyocytes (iPS-CMs) by synchronizing the beating of the cells to their own heart rate. Created via a close and continuing partnership between an academic laboratory and a science museum, *GHCB* employs human stem cell-derived tissue and demonstrates that stem cell technology can be used to create innovative educational experiences engaging to museum visitors. The exhibit was available to visitors in its current form from January to March 2020 at the Exploratorium, an interactive museum of science, art and human perception in San Francisco. Although the Exploratorium temporarily closed in March of 2020 due to the COVID-19 pandemic, the exhibit will remain part of the museum’s permanent collection upon reopening. We conducted visitor evaluation to gauge visitor interest and understanding and assess if the *GHCB* exhibit prompted visitors to think further about important scientific topics.

## Results and Discussion

We designed *Give Heart Cells a Beat* (*GHCB*) as a way to allow museum visitors to interact dynamically with cells in culture, (Fig 1a-b, Supplementary Video 1-2, Supplementary Figure 1). In *GHCB*, a handlebar-style heart rate sensor (Supplementary Figure 2a) is connected to a pacing electrode that is inserted in a cell culture plate containing iPS-CMs (Supplementary Figure 2b). When visitors place their hands on the handlebar, the electrode synchronizes the beating of the cells in culture to their heart rate in real-time. A live feed from the microscope is projected on a large screen for the visitors to observe, and an interpretive text overlay encourages them to interact dynamically, for instance by performing activities that increase or lower their heart rate (Supplementary Table 1). The microscope, with cells in an environmental control chamber, is inside the museum’s laboratory facility. It is visible through a large glass window to help visitors appreciate the scale and authenticity of the specimen^14^ (Supplementary Figure 2c).

**Figure 1.**
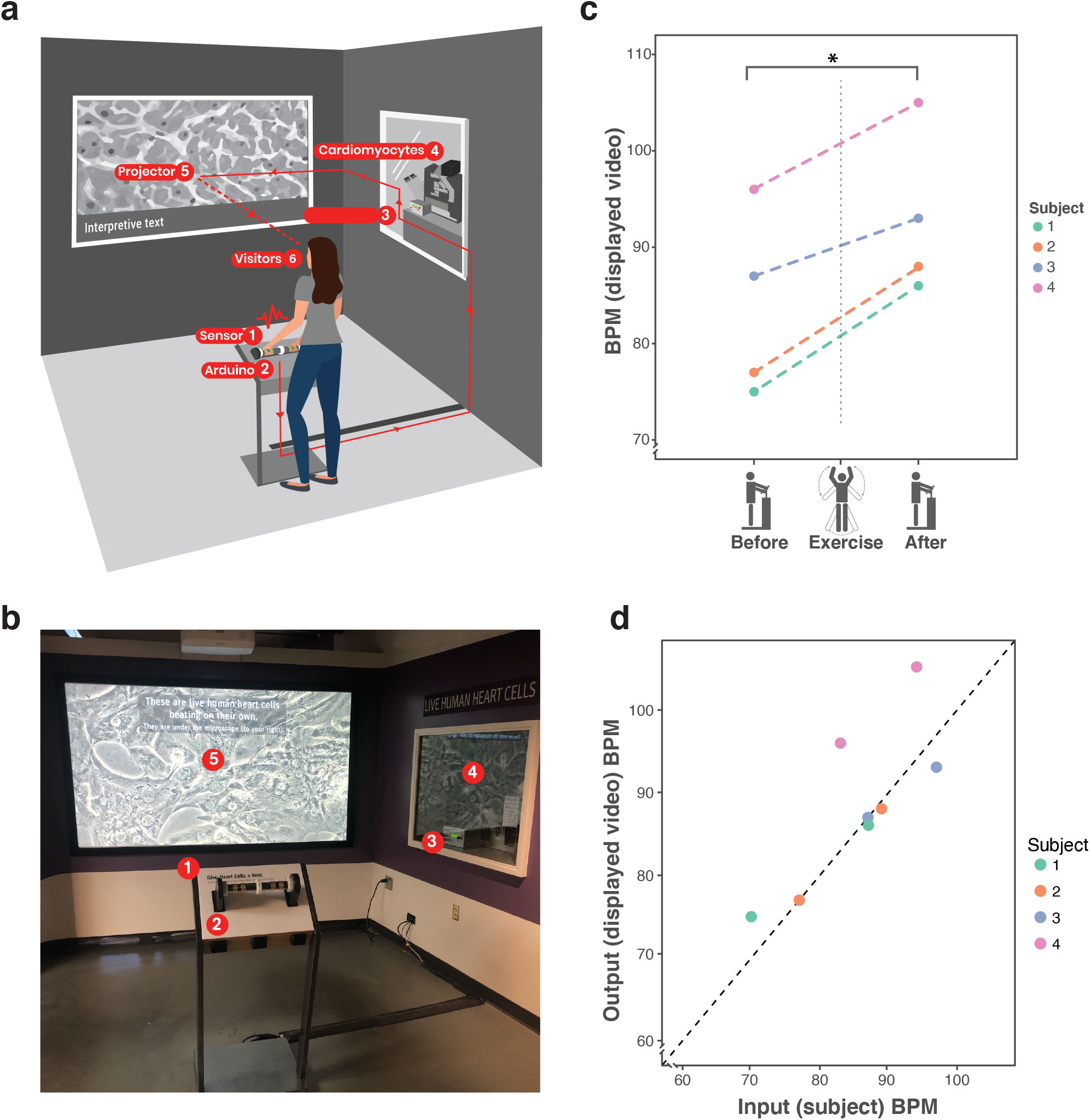
The Give Heart Cells a Beat exhibit enables visitors to accurately synchronize the beat rate of iPS-derived cardiomyocytes to their heart rate in real time. (**a**) Schematic diagram of exhibit’s functioning. The different parts of the exhibit (1-5) form a circuit that is ‘closed’ by the visitor (6). (**b**) Photography of actual exhibit layout. (**c**) The GHCB exhibit responds to changes in the heart rate of visitors. Displayed beat rate (obtained by video analysis) of iPS-CM cells in culture synchronized to four different users before and after performing light exercise (N=4, paired t-test p-value=0.004). (**d**) Comparison of actual heart rate of the user (measured by a medical-grade heart rate monitor) to the displayed beat rate for the cells in culture (measured by automated analysis of exhibit output video). Correlation is very good except for user 4, probably due to an improper adjustment of the heart rate monitor (linear regression, N=8, R^2^=0.97, p-value=0.0002).

To test the functioning principle of *GHCB*, we asked a set of volunteers to interact with the exhibit before and after performing a short exercise routine. In all cases, we were able to observe a significant increase in the displayed beat rate after exercise (Figure 1c). In addition, we observed a near perfect correlation between the actual heart rate of users and the beat rate of the cells in the projected video (Figure 1d). This allowed us to conclude that *GHCB* accurately and sensitively synchronizes the beat rate of iPS-CMs to the heart rate of the user in real time.

To assess visitors’ reactions to *GHCB*, we observed unprompted museum visitors as they interacted with the exhibit, then approached and interviewed a subset of those visitors (for detailed results and discussion, see Supplementary Information 2). The average time spent by the visitors in the exhibit was relatively long compared to other exhibits in the museum (Supplementary Information 2). GHCB’s “holding time”, a standard metric used to measure engagement^6,8,15^, suggests that visitors found the exhibit engaging. In addition, most of the visitors interviewed reported finding the exhibit interesting (Figure 2a), the two most frequently given reasons being interactivity and the opportunity to see live heart cells (e.g., “[I usually] don’t see heart cells because they are in you”; “[The exhibit] makes it feel very intimate and tangible”). The interviews also revealed that most visitors understood they were looking at heart cells or tissue, with a smaller majority reporting that these cells were of human origin (Figure 2b-c). In addition, 90% of visitors reported thinking more about their hearts, including health or a comparison to others (e.g., “I wondered about what condition my heart really is in, and it made me interested in taking more care of my heart.”), while 30% of visitors mentioned the technology behind the exhibit (e.g., “how did they do that? How much electricity can you use?”), and 20% talked about stem cells (e.g., “I was just interested in the fact that they were able to recreate human heart cells with stem cells”). Taken together, these findings suggest that *GHCB* provided an engaging, relatable experience and sparked thought about the heart and, to a lesser extent, advanced biology concepts such as stem cells or electrophysiology.

**Figure 2.**
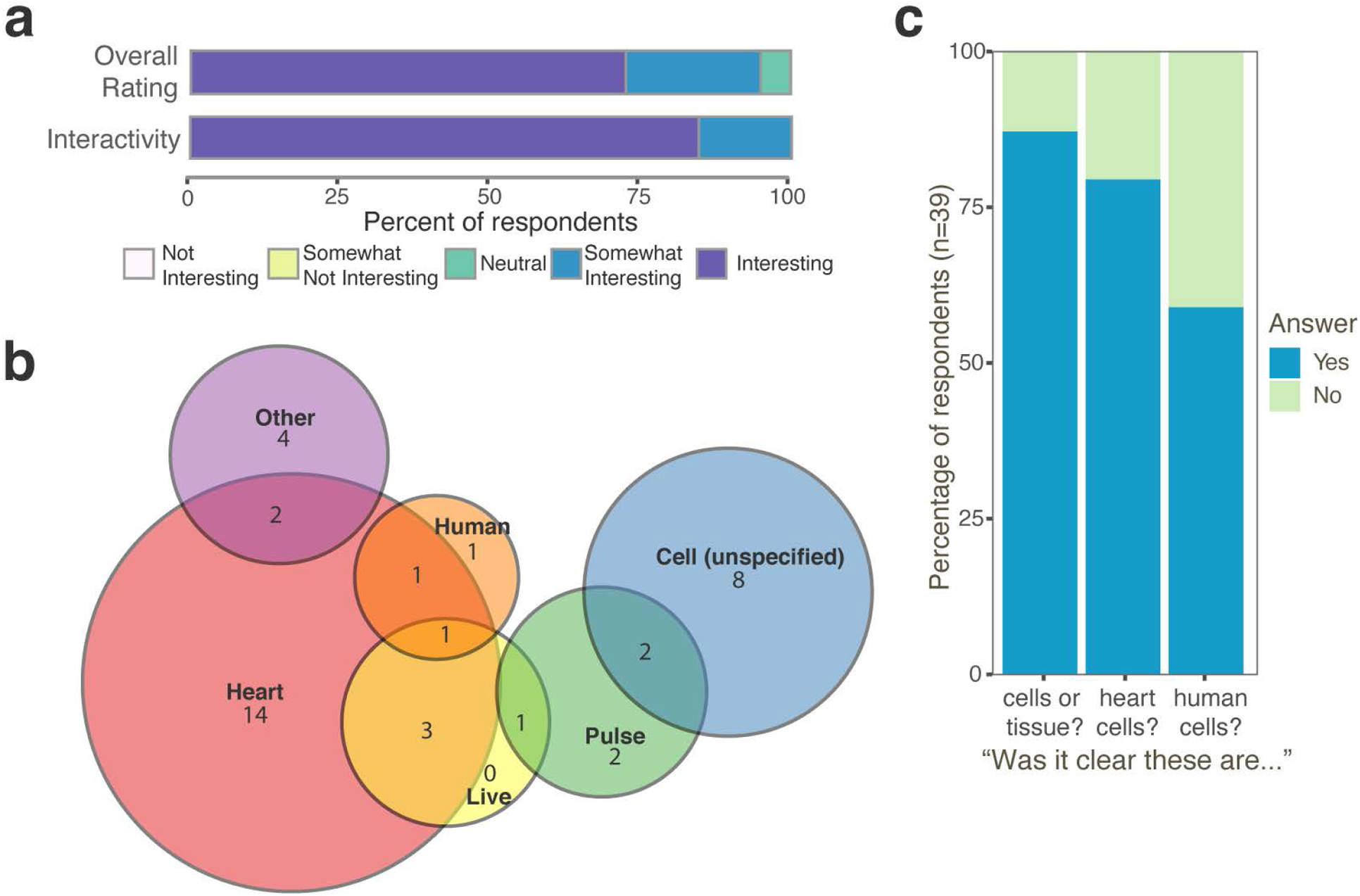
Visitor Evaluation. (**a**) Tally of interest ratings reported by visitors interviewed after using the exhibit for the overall exhibit (*n* = 40) and, specifically, its interactivity (n=39). (**b**) Venn Diagram of self-reported key terms visitors used to describe what they remembered seeing on the screen (*N* = 39). (**c**) Visitor responses when asked directly about what was shown in the screen.

Ultimately, GHCB is made possible by a close collaboration between a state-of-the-art academic stem cell research laboratory, which provides the cell specimen and scientific guidance, and a science museum which provides pedagogical and design expertise and access to a wide public audience (Figure 3). The iPS-CM cells are produced, differentiated, and stored in the academic lab as part as a routine protocol. Then, frozen vials are transferred to the museum laboratory to be thawed and purified for use. The post-mitotic nature of cardiomyocytes allows extended culturing, minimizing labor-intensive feeding or passaging steps and the use of consumable materials. In our experience, cells tolerate 4-5 months of intermittent pacing in the exhibit before becoming refractory to pacing, correlating with the onset of sarcomeric abnormalities (Supplementary Figure 3). Typically, leftover cells that were no longer of use to the academic lab were used, though if cells were made explicitly for the exhibit, a single differentiation batch could maintain the exhibit for several years (e.g. four 10cm dishes yielding ∼40 million pure iPS-CMs will allow the exhibit to run for a conservative estimate of 5 years, with an approximate cost of $500 in reagents and 10 hours of labor). Thus, cells can easily be made available to the museum, circumventing otherwise-prohibitive costs of purchase. In this case, the museum leveraged an existing laboratory (with a basic cell culture facility), microscopy facility, and museum staff with training in cell culture to support the exhibit. In return, the academic lab benefits from the increased understanding and appreciation of their research by a broad audience, while the museum bolsters its mission by offering a completely unique experience to its visitors, through communicating cutting-edge scientific and research concepts to the general public.

**Figure 3.**
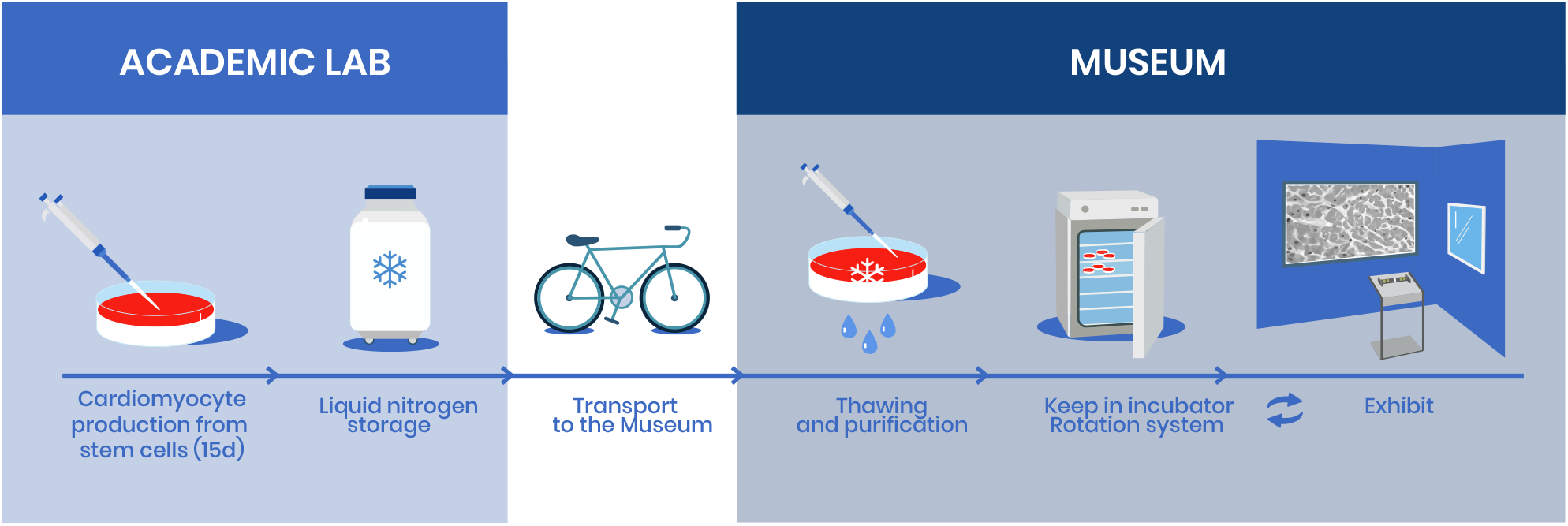
Give Heart Cells a Beat is an interactive, permanent exhibit enabled by direct collaboration between an academic lab and a museum. Stem cell-derived cardiomyocytes are produced in an academic lab (normally leftover from other experiments) and frozen down. When requested, frozen vials are handed over to the museum, where they are thawed and lactate purified (this step can instead be done at the academic lab). Cells are kept in an incubator in the museum and rotated in and out of the exhibit to prevent excessive deterioration.

The *Give Heart Cells a Beat* exhibit is a novel educational implementation of stem cell technology and utilizes the uniquely dynamic behavior (beating) of cardiomyocytes to enable a real-time interactive experience. The exhibit is a proof-of-concept of the potential for stem cell technology to impact the field of science education, and a pioneering example of the application of biotechnological advances to provide improved tools for science communication. The *GHCB* exhibit was conceived in 2015 by Exploratorium staff and Gladstone Institutes scientists, and builds upon an existing collaboration which featured mouse stem cell-derived cardiomyocytes. Currently, *GHCB* enables a continued exchange of reagents and information between the institutions, allowing for constant refinement of the exhibit and the protocols, and a foundation for further interactive exhibit development featuring stem cell-derived cell types such as neurons, blood vessels, or immune cells, or other state-of-the-art biological research tools and advances.

In this study we have reported the development of *Give Heart Cells a Beat*, the first science museum exhibit to allow dynamic, real-time interaction between museum visitors and cultured human stem-cell derived cells. Our evaluation study with visitors highlights the importance of interactive design to foster engagement with biology content, and also suggests that the use of heart cells provides a relatable hook for visitors that may help bridge to further concepts including cardiology, stem cells, or electrophysiology.

This exhibit is a product of an effective and sustained collaboration (>15 years) between academia and a science education center and an example of how long-term collaborations can produce novel implementations of current technology to communicate scientific advances to the public. As trusted public institutions^16,17^ museums and science education centers are uniquely positioned to collaborate with research labs in raising awareness of the importance of biomedical research, particularly in light of the COVID-19 pandemic. Exhibits like *GHCB* can contribute to this task, particularly as the cell line used in the exhibit has been used in published research on the cardiac impact of COVID-19^18^. The *Give Heart Cells a Beat* will remain a permanent exhibit at the Exploratorium museum in San Francisco, where thousands of visitors have already interacted with it.

## Methods

### Exhibit Design

Upon approaching the exhibit, the visitor triggers a proximity sensor, prompting an on-screen graphic to appear with instructions to grab a hand-activated heart rate sensor (Insta-Pulse). The visitor’s heartbeat is detected by the handheld sensor and sent via an Arduino device to a Myopacer Cell Stimulator (Ionoptix), which then delivers a pulse (biphasic, 10v, 10ms duration) to a plate of iPS-CMs via a submerged 2-prong carbon electrode (IonOptix). The cardiomyocytes contract in response to each detected beat (supplementary figure xxx). A camera captures the beating of the cells and routes it through a program which overlays text instructions to a projector for the visitor to view. As the visitor continues to use the exhibit, the on-screen instructions change, prompting different activities. The Arduino also sends the signal to a haptic feedback system (Uxcell), which send a vibration pulse to the handlebar in response to each beat, helping visitors monitor their own heart rate and verify the synchronization of the cells. The microscope system (Zeiss Axiovert 200M) is equipped with an environmental control chamber enabling cells to be kept on exhibit for multiple days.

### Human iPSC culture and differentiation into cardiomyocytes

Cells from the WTC human iPSc cell line, derived from a healthy male subject^19^ were maintained in mTeSR1 (STEMCELL Technologies) media on growth-factor-reduced Matrigel (8 μg/ml, BD Biosciences) and passaged every 3–4 days using Accutase (STEMCELL Technologies). ROCK inhibitor Y-27632 (10 μM, Selleckchem) was added to the media for 24 h after each passage. Cells were differentiated into cardiomyocytes as described previously^19^. Briefly, iPS cell cultures were given 12uM CHIR99021 (Tocris) in RPMI 1640 (Gibco) with 2% B-27 supplement without insulin (Gibco) approximately 72 hours after plating (day 0). Media was changed to RPMI/B27 without insulin a day later, and then RPMI/B27 (without insulin) containing 5 μM IWP2 (Tocris). After another 48 hours, the media was changed to RPMI/B27 containing insulin. Fresh RPMI/B27 was exchanged every 3–4 days thereafter. On day 15, cells were harvested using 0.25% Trypsin (Gibco) and either replated for lactate purification (see below) or frozen in CryoStor media (BioLife Solutions). Cells were stored in Liquid Nitrogen tanks until transferred to museum facilities for thawing.

### Cell Thawing, Purification and Maintenance on Exhibit

Frozen vials of differentiated cells were transferred from the Gladstone Institutes to the Exploratorium on dry ice. Cells were thawed in 6-well plates and, if they had not been purified, cardiomyocytes were enriched using a metabolic selection method^20^. Briefly, three days after plating, media was replaced with DMEM without glucose (Gibco) supplemented with 4mM lactate (Sigma). Lactate media was exchanged every other day to a total of 3 times. Cells were then maintained in RPMI/B27 with 0.5% Penicillin-Streptomycin (ThermoFisher) or Antibiotic-Antimycotic (ThermoFisher) in Matrigel-coated dishes.

### Pacing on Exhibit

Once purified, cardiomyocytes were kept in an incubator for a variable length of time, until the autonomous beating rate is around 30-40 beats per minute. When necessary, this was assisted by the use of Ivabradine (Sigma Aldrich) or by decreasing the temperature of the cultures. Each plate of cells was paced on the exhibit for 2 days in media containing an antioxidant, L-ascorbic acid (212.5 ug/mL, Sigma Aldrich), then allowed to recover 1-2 weeks without pacing before being used again. By rotating through a group of 6-10 plates in this manner, oxidative stress was minimized and cell cultures remained viable for use for multiple months.

### Beat rate analysis

Randomly selected volunteers were asked to interact with the GHCB exhibit before and after performing exercise (15 side-straddle hops, ‘jumping jacks’) while wearing a Kardia (AliveCor) heart rate monitor for actual heart rate tracking. For automated analysis of beat rate of video output, video clips recorded from the exhibit screen were analyzed using the Pulse Video Analysis platform^22^ (Dana Solutions).

### Visitors’ evaluation

A total of 62 randomly chosen museum visitors were observed, of which 40 were interviewed. For a detailed discussion on methodology and results, see Supplementary Information 2 - Visitor Research and Evaluation for *Give Heart Cells a Beat* exhibit. For the analysis of key descriptors in the visitor interviews, comments were manually annotated and classified. The term “cells-unspecified” refers to answers that acknowledged seeing cells without any other descriptor, while “other” was used for answers that didn’t fit in previous categories (“an image”, “blood”, or simply “weird stuff”). Data for Likert scores and key terms was plotted using R (version 3.5.3).

### Immunofluorescent staining of sarcomeres for qualitative evaluation of cell health

Cell plates from the exhibit were fixed in 4% paraformaldehyde for 15 minutes at room temperature. They were then washed with PBS containing 0.1% Triton X-100 (PBS-T), then blocked in 5% bovine serum albumin (BSA, Sigma-Aldrich) in PBS-T at room temperature for 1 hour. For sarcomere imaging, Alpha actinin antibody (A7732, Sigma-Aldrich) was then diluted in 5% BSA solution and incubated overnight at 4°C. Cells were washed in PBS-T and then incubated with secondary antibody (Alexa Fluor 594 goat anti-mouse IgG; Molecular Probes) diluted in 5% BSA solution for 1 hour at room temperature. Nuclei were stained using DAPI (Vector Laboratories) and cells were imaged using a BZ-X700 microscope (Keyence).

## Supporting information

Supplementary Information 2 - VRE results and discussion

## Acknowledgements and funding

*Give Heart Cells a Beat* is an exhibit made possible through generous support of the Gordon and Betty Moore Foundation, The Troy and Leslie Daniels Fund for Life Sciences, and Genentech. J.P.B. received funding from *La Caixa* foundation and the American Heart Association during the span of the work described in this manuscript.

B.R.C. receives funding from the NIH and the Gladstone Institutes.

The authors would like to thank Alexandre Ribeiro for his advice in the conception of this exhibit, Angela Armendariz, Ray Larsen, Kevin Boyd, Matt Trocker, Dana Carrison-Stone, Veronica Johnson and members of the Exploratorium Living Systems department for their help with the setup and maintenance of GHCB, and Rosario Sotelo, Joanna Steinhardt, and Rodney Wilson for collecting the visitor data. We also thank the Gladstone Stem Cell Core for their support and experimental expertise, and the Gladstone Editorial team for constructive feedback on this manuscript.

Figures 1a and 3 were designed and generously provided by Paula Marengo (www.marengocreative.com).

## Author contributions

J.A.P.-B., S.J.R. and J.M. designed the study, performed the experiments and analyzed the data. J.A.P.-B. and S.J.R. performed cell culture, differentiation, and maintenance of cells. C.C. designed the *GHCB* exhibit. B.R.C. and K.Y. supervised the study. All authors contributed to writing the manuscript and figure preparation.

