## Supplementary Information 1 - Supplementary Figures and Tables for "Synchronization of iPS-derived cardiomyocytes to visitor heartbeat in an interactive museum exhibit"

**Supplementary Table 1** - Interpretive text showed on screen to guide visitor experience on GHCB

| Time | Text |
|---|---|
| Before approach | "These are live human heart cells beating on their own. They are under the microscope (to your right)." |
| When near | "The handlebar senses your heart rate and sends it to the live heart cells under the microscope. [image] Grasp the handlebar." |
| Countdown, once the visitor grabs the sensor. | "The handlebar senses your heart rate and sends it to the live heart cells under the microscope." |
| In the interaction (each screen is shown for 10 seconds, with a 2-second interval between screens). | (1) "These human heart cells were grown from stem cells in a lab."<br>(2) "How do these heart cells respond after you do some exercise?"<br>(3) "How do they respond to your friend's heart beat?"<br>(4) "Can you slow down these cells' beating?"<br>(5) "Look at the microscope setup that keeps these cells alive." |

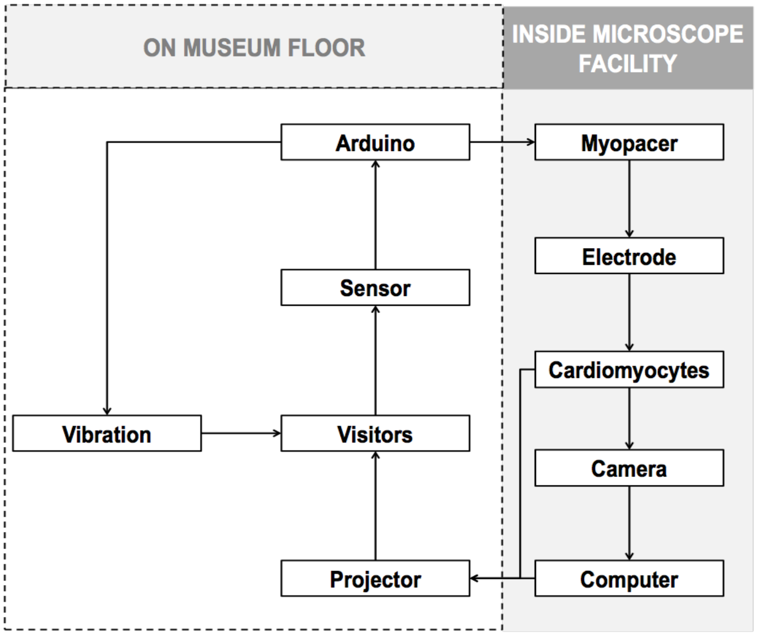


**Supplementary Figure 1** - Exhibit design diagram. The visitor's heartbeat activates the pacing of the cardiomyocytes in culture. Visitors receive input in the form of the projected video and the pulses to the handlebar.

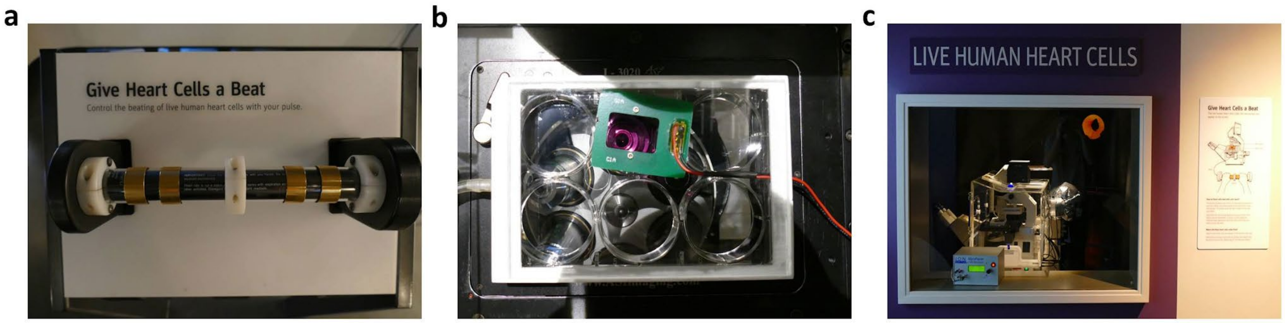


**Supplementary Figure 2** - Detail of exhibit parts. a) Hand-held heart rate sensor used by visitors to interact with the exhibit. The handlebar is equipped with a vibration device that vibrates with the perceived beat rate. b) Close-up of the plate of cells in culture, with the Myopacer electrode (green plaque, orange wire) inserted. c) A visitor's view of the microscope and environment chamber. A wall graphic (right) and large type letters (above) explain the content to visitors. The pacing device is also on display (bottom).

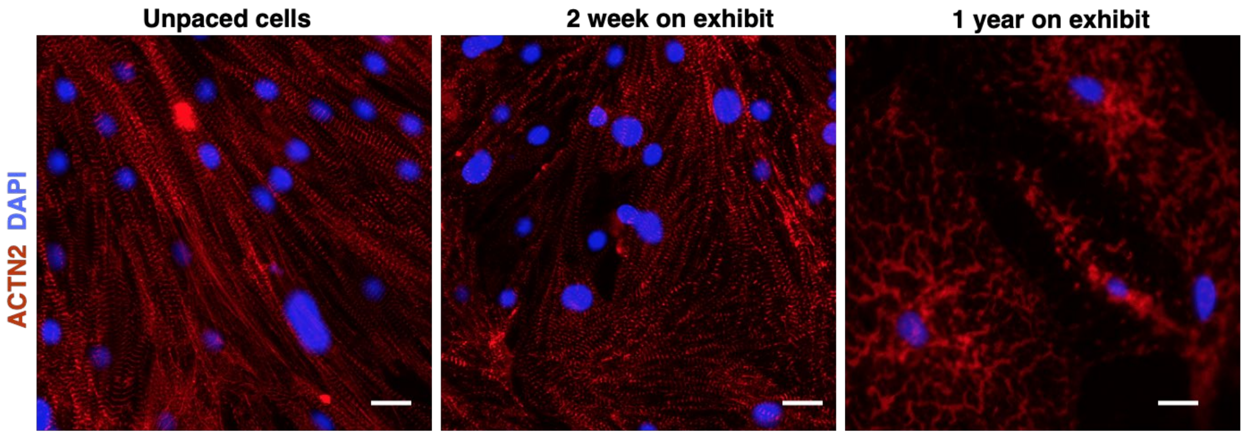


**Supplementary Figure 3**. Functional cardiomyocytes develop sarcomeric abnormalities under prolonged pacing periods. Sample images of cultures of cells that had been on exhibit rotation for varying amounts of time (2 days per week). Unpaced cells (left) show high confluence and display well defined sarcomeres, with few sarcomeric material aggregates. After 2 weeks in the exhibit (center), cells show longer and more defined sarcomeres, with an increased amount of sarcomeric material aggregates. Cells that have cycled through the exhibit for much longer (1 year, right) are very sparse (probably due to cell death) and much larger, with large protein aggregates all over the cytoplasm instead of distinguishable sarcomeres. (scale bar = 10µm)

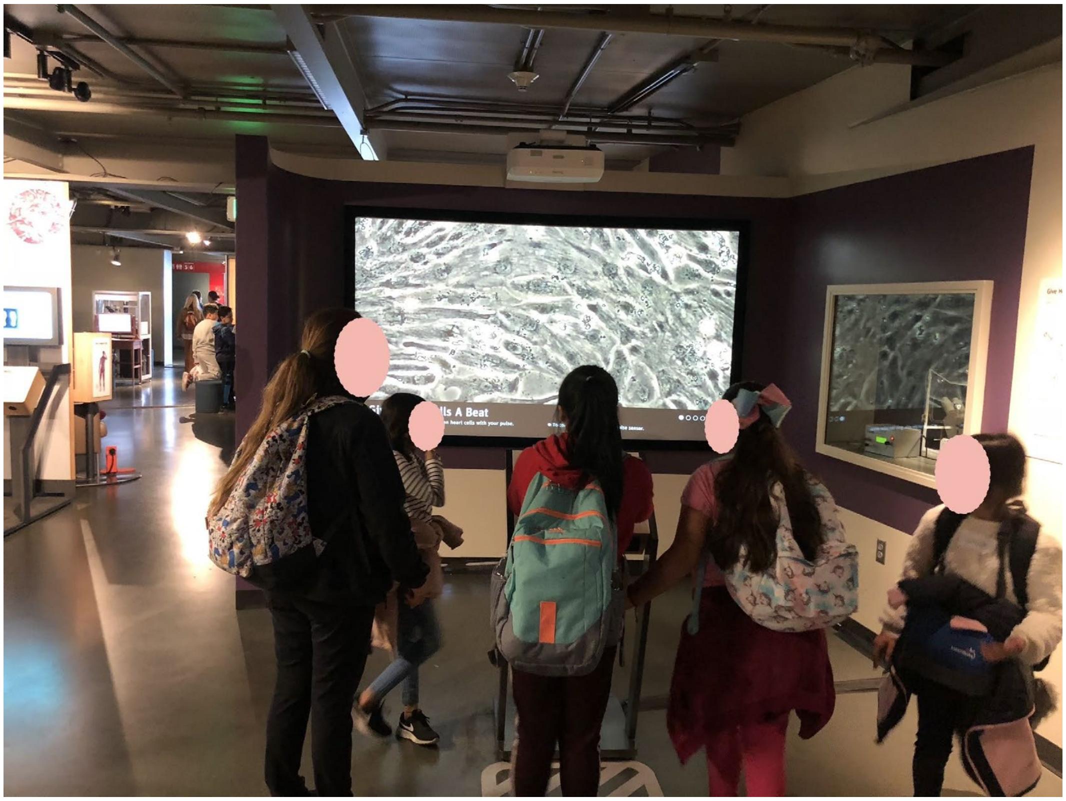

**Supplementary Figure 4**. Visitors using the exhibit, prior to the finalization of graphics referred to in this article

**Supplementary Video 1 - Example of projected cells beating.**

Note: The on-screen interpretive text has changed since this video was recorded. Current on-screen text can be found in Supplementary Table 1.
