## Supplementary Information 2 - VRE results and discussion for "Synchronization of iPS-derived cardiomyocytes to visitor heartbeat in an interactive museum exhibit"

### Supplementary Information 2 - Visitor Research and Evaluation results and discussion

##### Purpose

An evaluation was conducted to determine what, if anything, visitors found engaging and worthwhile about GHCB. More specifically, it looked for evidence for and against a subset of the exhibit's design assumptions, summarized here:

- GHCB provides a valuable visitor opportunity to see and interact with a microscopic, in vitro sample, in real-time.
- The use of human heart muscle cells as a sample provides a compelling, immediately relatable hook for visitors.
- This hook provides a bridge to other, more difficult concepts like stem cells, electrophysiology, and cardiology.

##### Method

The study collected data over four weekend days and one holiday in the winter of 2020. An evaluator stood near the exhibit and observed every third visitor who (a) appeared 8 years old or older and (b) stopped in front of the exhibit's handlebar for more than ten seconds. When that individual left the exhibit area, the evaluator approached that visitor for an interview (for details on the questions see Supplementary Information 3 - VRE Form). If that individual was a child or teenager, they asked for consent from the accompanying adult to interview the minor. In total, the evaluation study observed 62 and interviewed 40 visitors, of which one dropped out in the middle of the interview. Most participants chosen through random systematic selection in this study were adults (See Sup Table VRE.1 below).

**Sup Table VRE.1 - Demographic information for evaluation study participants**

| | Count Observed (percentage of 62) | Count Interviewed (percentage of 40) |
|---|---|---|
| Age Group | | |
| Child | 8 (13%) | 6 (15%) |
| Teen | 4 (6%) | 3 (8%) |
| Adult | 50 (81%) | 31 (78%) |
| Gender | | |
| Female | 33 (53%) | 18 (45%) |
| Male | 29 (47%) | 22 (55%) |

**Interview Questions**

1. How interesting did you find that exhibit? Would you say that was …

| Not Interesting | Somewhat Not Interesting | Neutral | Somewhat Interesting | Interesting |
|---|---|---|---|---|

   a. What made it ___________ for you?

2. Did you find out anything new at the exhibit?

3. Is there anything that you're wondering or became interested in after using this exhibit?

4. We're wondering how easy it was to understand what's being shown. Without looking back at the screen, do you remember what was on the large screen? What was it showing?
   a. Was it clear that these are cells or tissue? YES NO
   b. Was it clear what kind of cells these are? YES NO
      i. [Clarify HEART] So, was it clear that these are heart cells? YES NO
   c. Was it clear that these are human cells? YES NO
      i. Does knowing that they are human cells make it….

| Less interesting | More interesting | Doesn't really make a difference |
|---|---|---|

      ii. How so? Why's that?
   d. Do you think what you are seeing here are real? Or, a simulation? Or a canned video?
      i. What makes you think this?

5. Were you able to get these to beat with your own heart beat? YES NO

   [If YES in any way]
   a. Was that

| Not Interesting | Somewhat Not Interesting | Neutral | Somewhat Interesting | Interesting |
|---|---|---|---|---|

   b. What made that _______?

6. Did the exhibit make you think about your own heart? [Probe: How so?]

7. Do you have any special interest or background that might have helped you understand what you saw, perhaps from school or a hobby at home?

**Results and Discussion**

As part of the evaluation observation, we collected holding time data, the amount of time the subjects stayed at the exhibit. Holding time is a well-established metric of engagement in the museum field where visitors themselves decide how long to attend to an exhibit[1]. The median holding time for GHCB was one minute (Sup Fig VRE.1). As a point of comparison, visitors spent approximately ten seconds to a little under two minutes per exhibit in an earlier Exploratorium life sciences collection, with the median holding time being 42 seconds averaged over 37 exhibits[2]. The long holding times suggest that visitors were engaged with GHCB, since a disinterested visitor could easily have left for another, more exciting exhibit nearby.

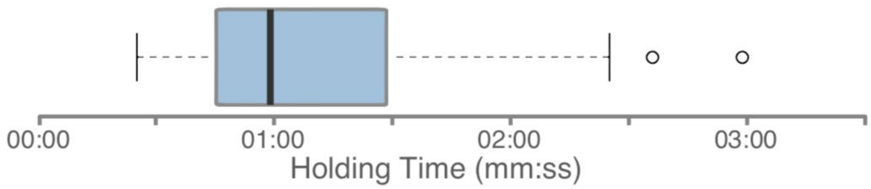


**Supplementary Figure VRE.1 -** Distribution of holding times for visitors of the Give Heart Cells a Beat exhibit (n=62).

Most (29/40) of the visitors we interviewed rated the exhibit as *Interesting* (5), with 95% of them (38/40) rating it above *Neutral (3)* on a five-point Likert scale from *Not Interesting* (1) to *Interesting* (5) (main text Figure 2a). Further probing provided insights into what visitors found interesting; the two most frequently given reasons were:

- They liked the interaction, particularly the ability to synchronize their heartbeat to the exhibit (19/40). For example,

  *V11: Seeing it follow our heartbeat. Cool beans.*

  *V20: The fact that it synchronizes to your heart beat [was interesting]*

  *V37: The idea that you can be synched with cells themselves. [I] haven't seen anything like this before.*

- Visitors liked seeing the heart cells (15/40).

  *V1: [I usually ] don't see heart cells because they are in you. [The exhibit] makes it feel very intimate and tangible.*

  *V28: [It's interesting] to see what's happening in my body, visually.*

  *V58: It shows what your cells are doing while heart is beating*

These responses give initial support to our original assumption that GHCB can provide a valuable opportunity for visitors to see and interact with an in vitro, microscopic specimen.

To unpack how the exhibit's interactivity contributed to visitors' engagement, we asked the study subjects to rate how interesting it was to synchronize their heartbeats and why that was interesting (or not). On a 5-point scale, most (33/39) visitors reported that the ability to interact with the heart shown on the large screen was *Interesting* (5), with the remainder rating the interactivity as *Somewhat Interesting* (4) (main text Figure 2a). They gave a variety of reasons for their ratings, the two most prevalent, with at least 25% respondents, being that:

- It allowed them to visualize how their hearts may be beating (14/39). For example,

  *V7: We know we can check our own pulse but to see it reacting is pretty cool.*

  *V43: Just cause seeing my cells move to my actual heart beating was cool.*

  *V56: You can't see how it is every day because you can't just look at yourself you need something that can see stuff really small*

- The ability to control cells was impressive to visitors (9/39). For example,

  *V3: The fact it can synch up. [I] didn't even think it was possible.*

  *V17: The fact that we can control [other] cells with our own pulse.*

The interactivity, therefore, seemed to allow a majority of visitors to see and think about how their own hearts may be beating that goes beyond simply controlling cells.

A large part of allowing visitors to make that connection between the cells on the screen and their own hearts lies in understanding the projected image in front of them. To gauge if visitors knew they were looking at live human heart cells, we asked them to describe what they remember seeing on the screen and coded their responses. Table VRE.2 tallies what visitors called out, while Figure 2 in the main text shows the Venn diagram for visitors' descriptions about the type of cells they thought they saw. We note that only one person self-reported seeing "live" + "human" + "heart" cells.

Because visitors' self-reports may incompletely capture what they thought, we also asked them directly if they knew what they saw on the screen were live, human heart cells. When asked, most visitors reported thinking that they were looking at cells or tissue (34/39), and many knew they were looking at heart cells (31/39), with a smaller majority (23/39) reporting thinking that they were human cells or tissue (Figure 2b-c in main text).

**Sup Table VRE.2 - Tally of what visitors thought was shown on the screen.**

| What visitors thought was on the screen | Count (percentage of 39) |
|---|---|
| Heart Cells | 21 (54%) |
| Cells (unspecific) | 11 (28%) |
| Other (i.e., stem cells, blood, weird stuff) | 6 (15%) |
| Pulse | 5 (13%) |
| Live Cells | 5 (13%) |
| Human Cells | 3 (8%) |

Although we had assumed that using human heart cells would provide an immediately relatable hook for visitors, we found that visitors did not readily interpret what they saw as being human, despite the annotations on the screen. We think that this could be because the *human* nature of the cells was not obvious from the projected image (in contrast to *cells*, which have a more familiar organic look or *heart*, which is more readily identified by their beating motion).

When they were informed during their interviews that these were human heart cells, a majority of visitors thought that using human cells made the exhibit experience more interesting. More specifically, of these 22 visitors, a majority (15) thought that doing so made the exhibit experience more relatable. For example,

*V5: It's us.*

*V17: Because you can relate to it. They're inside you, in your heart.*

*V53: (laughs) It personalizes it.*

Alternatively, we were surprised to find that a large minority (17/39) of visitors felt that having human cells did not really add to the exhibit. We speculate that this could be due to a number of factors, including but not limited to: (a) it is difficult for some visitors to relate to cells in a culture plate; or (b) there is a lack of understanding of the differences between human and animal cells and the difficulties of using human cells. As some of these visitors explained:

*V11: It's still a heart cell.*

*V60: Any living creature's heart cell [is the] same to me.*

Independent of this, at the end of their exhibit experience most (35/39) visitors reported thinking more about an aspect of their own hearts. These included thinking about the health and condition of their own hearts (16/39); for example

*V4: I'm interested in heart because we have heart disease in our family so makes me think about that.*

*V28: I wondered about what condition my heart really is in, and it made me interested in taking more care of my heart.*

*V33: Yeah, you should take care of your heart.*

Some (14/39) visitors also reported thinking about how their hearts look and behave; for example

*V9: Yes, is this regular? Is this what it is supposed to look like?*

*V21: if the heart really looks that way.*

*V43: Yeah, going back to how they all look like. Does mine look like this inside?*

A few (3/39) visitor thought about how their heart compares to others; for example

*V37: My heartbeat is a little slower than his. He's a runner, so to compare them.*

These findings suggest that using human heart cells can be a promising way of making the interactive experience relatable for some but not all visitors, and their effectiveness, in turn, depends in part in clearly conveying what they are.

One of our design assumptions was that using live human heart cells can help incite curiosity and provide a starting point to exploring more difficult concepts such as stem cells, electrophysiology, and cardiology. When they were asked during their interviews what they became curious about, some (12/39) visitors talked about the technology behind the exhibit; for example:

> *V1 : And also how did they do that? How much electricity can you use without killing the cells? the electricity from your heart to the dish.*

> *V35: I see what's happening I would like to learn how it synchs up to my pulse rate.*

The exhibit piqued other (8/39) visitors' interest in stem cells:

> *V22: Just how they [the cells] were generated.*

> *V28: I was just interested in the fact that they were able to recreate human heart cells with stem cells.*

In addition, other visitors became curious about heart cells in general (7/39) and in their heart in particular (5/39). In contrast, a large minority (17/39) reported not becoming more curious about anything in particular after using GHCB.

**Conclusion**

In this short report we have summarized the results of an evaluation of the visitor experiences after interacting with the GHCB exhibit, as captured through in-person interviews. We found that the visitors valued very highly the ability to observe and, especially, interact with the microscopic specimen. We also found that, for a majority of visitors, the exhibit was thought-provoking and prompted curiosity about the nature of the stem cells used, the technology that makes the exhibit possible, and/or different facets of heart physiology and health. To our surprise, the interviews also revealed that a large minority of users did not feel that the *human* nature of the cells used added to their experience, pointing at a difficulty in conveying this aspect of the exhibit. Overall, the findings of this study indicate that the use of human stem cell derived cells in an interactive setting shows promise in engaging the visitor and sparking further explorations, but that the current exhibit will require further work and iterations to realize its full potential
